# Tracking the long way around: seasonal migration strategies, detours and spatial bottlenecks in Common Cranes wintering in Western India

**DOI:** 10.1101/2025.03.26.645416

**Authors:** Harindra L. Baraiya, Gaurav Sirola, Anju Baroth, R. Suresh Kumar

## Abstract

**Background:** The Common Crane *Grus grus,* being widely distributed, abundant, and relatively easy to monitor, with long-distance migrations across diverse and sensitive habitats, is well-suited as a sentinel species for assessing the ecological integrity of the Central Asian Flyway (CAF). While migration routes and stopover sites of the Common Crane in the CAF are documented, seasonal variations in migration strategies, potential detours, bottlenecks, and stopover habitat selection - important factors for flyway conservation planning - remain poorly understood. To address this, we tracked five individuals from Gujarat, western India, to breeding sites in southwestern Siberia using solar-powered GPS-GSM transmitters, enabling high-resolution, multi-season analysis of their migration strategies, routes, timing, detours, bottlenecks, and stopover habitat selection.

**Results:** The results highlighted seasonally differential migration strategies in Common Cranes. The analyses confirmed significant seasonal differences: Spring migration covered greater distances, lasted longer, and was slower, with higher stopover duration and reduced straightness. The observed migration routes deviated significantly from the simulated straight-line path, forming distinct detours likely to avoid the Hindu Kush mountains and take advantage of resource-rich areas. These detours resulted in routes that were 23.3% longer in autumn and 36.8% longer in spring compared to the shortest possible path. The tagged Common Cranes followed distinct seasonal migration routes that converged in Turkmenistan, creating migratory bottlenecks in Southern Turkmenistan, Afghanistan and Pakistan. At migration stopover sites, Common Cranes showed the highest probability of habitat use in bare ground (0.18) and cropland (0.13), suggesting a preference for open landscapes by the species.

**Conclusion:** Our findings reveal the adaptive strategies of Common Cranes in response to seasonal and topographical challenges. Cranes undertake substantial detours to optimize energy expenditure and access favorable stopover habitats. The identification of migratory bottlenecks in Turkmenistan, Afghanistan and Pakistan highlights the region’s critical importance for conservation efforts for migratory birds.

## Background

Avian migration is a periodic, repetitive, and large-scale movement, taking place at the same season each year, with individuals commuting between breeding and non-breeding areas (1–3). Migratory species are exposed to the highest risk during their migratory journeys (4,5) encountering a spectrum of natural and anthropogenic challenges during migration. These challenges include extreme weather changes, habitat alterations, hunting and poisoning along with many other (6–8). Certain avian migratory species allocate more than half of their annual life cycle to traveling between breeding and wintering habitats (9). Adversities encountered during these transitions significantly contribute to a considerable proportion of annual mortality across various species (4,10). Considering these, extensive studies have been undertaken to understand the patterns, drivers, and consequences of migratory movements (11–13), including the evolution of partial migration (14), stop-over site ecology (15–18), and the influence of environmental changes on the shifting migratory timings and movements (19–23).

A flyway concept provides a spatial framework for the management and conservation of migratory avian species across political boundaries as it encompasses the multiple migratory routes spanning countries along the entire geographic range (24). One such flyway is the Central Asian Flyway (hereafter, CAF) spread across a large continental area of Eurasia between the Arctic and Indian Oceans and the associated island chains (25). This flyway is located in the Asia-Pacific region where the highest number of threatened soaring bird species have been reported (6). However, this is also the flyway that remains comparatively less studied in terms of migratory ecology and conservation research (26). The Asian Flyways are experiencing large-scale threats, including hunting migrant birds (26–28), extensive habitat loss and urbanization, and the limited effectiveness of protected area networks in preventing habitat degradation-factors likely to cause population-level impacts on migratory bird species (29,30). Understanding a species movements and migratory connectivity can help understand the species-specific and route-specific threats including the spread of infectious diseases that may have population-level consequences (31,32).

Among the numerous migratory species that travel along the CAF, the Common Crane *Grus grus* is one of the widespread species that winters in tens of thousands in India. Although the latest IOC World Bird List (33) does not currently recognize any subspecies for *Grus grus,* morphological and genetic studies have proposed four subspecies: the Western Common Crane G. g. *grus* in Europe; the Transcaucasian subspecies G. g. *archibaldi* in the Inner Caucasus/borderland of Turkey, Georgia, Armenia and Iran (34,35); the Tibetan subspecies G. g. *korelovi* in Tibet in China, Kyrgyzstan, and Kazakhstan (36); and the Eastern Common Crane G. g. *lilfordi* in Asia from the Ural Mountains eastward to Northeastern Siberia (37,38). The Western and Eastern Common Cranes are geographically separated by the Ural Mountains (39). The population size of the Eastern Common Crane is estimated to be 125,000-130,000 individuals (40), of which more than 100,000 migrate from Western Siberia and Kazakhstan to the wintering grounds in India (41–43) and Central Asia in the Amu Darya River Valley (44–46).

The arid plains in the States of Gujarat and Rajasthan in Western India support large populations of the Common Crane that arrive from different parts of their breeding range (41–43). Few of the wintering sites witness huge congregations of Common Cranes and these are often species-specific flocks numbering in thousands (HLB unpublished count data). Previous studies on the migratory behaviour of the Common Crane population wintering in Western India have provided important, yet fragmented, insights into their movement patterns along the Central Asian Flyway. The first attempt to document the Common Crane migration routes was made by (41), who tracked two individuals from Rajasthan, North-Western India, to southwestern Siberia. A subsequent study by (42) expanded on this by tracking three additional Common Cranes from Gujarat, Western India, to Siberia. However, both studies were constrained to spring migration data only, primarily due to technological limitations such as the low battery capacity of earlier satellite transmitters. This necessitated a trade-off between recording frequency and device longevity, resulting in limited data collection across the full annual cycle. (43) later employed improved tracking technology to follow two Common Cranes from the Saurashtra region of Gujarat, documenting detailed migratory routes during both Autumn and Spring migrations, as well as movement on wintering grounds. Nevertheless, these prior studies were limited by the use of single-season or single year data, which restricted their ability to characterize migration patterns comprehensively across seasons or to delineate primary migration corridors and critical stopover sites.

Moreover, none of the previous studies on the Common Crane from the region systematically quantified two critical parameters in migration ecology-migratory detours and bottlenecks-despite their central role in shaping energy budgets, stopover strategies, and conservation priorities. Detours from the shortest possible migration route can reflect complex interactions between topographical barriers, atmospheric conditions, and the distribution of key resources, and are increasingly recognized as adaptive responses to migration constraints (47–49). Similarly, bottlenecks-areas where multiple migration trajectories converge due to environmental or geographic constraints-represent critical junctures where populations are most vulnerable to localized threats (50,51). Quantifying these parameters is essential not only for understanding the underlying drivers of route selection but also for identifying conservation priorities at a flyway scale.

To address these limitations, our study applied state-of-the-art solar-powered GPS-GSM transmitters to track five Common Cranes from Gujarat across two years, capturing fine-scale, seasonal migratory movements from Western India to the steppes in Russia and Kazakhstan. This dataset enabled an in-depth characterization of key migration parameters-including total migration distance, duration, speed, straightness and stopover duration-allowing us to analyze and compare Spring and Autumn migrations. Further, we investigated migratory detours and identified bottlenecks along the migratory route, mapped high-use areas, and assessed habitat selection at stopover sites. Together, these parameters help characterize seasonal migration strategies, providing insights into how movement patterns may be shaped by ecological factors such as pre-breeding refueling needs or the availability of favorable flight conditions. These findings provide a foundation for identifying conservation priorities and informing flyway-scale management efforts along the CAF.

## Methods

### Study population and GPS tracking

During the winter seasons of 2019-20 and 2021-22, efforts to tag Common Cranes were concentrated in Nal Sarovar Bird Sanctuary (22.8183 N; 72.0439 E, area: 12,000 ha) and Thal Lake Wildlife Sanctuary (23.1339 N; 72.4001 E, area: 699 ha) in the Western Indian state of Gujarat. Both sanctuaries are designated as Ramsar Sites under the Ramsar Convention and harbour a substantial population of the Common Crane, estimated to be as high as 30,000 (based on unpublished data from regular, systematic roost counts by H LB).

We tagged five adult Common Cranes with solar-powered GPS-GSM leg-mount transmitters (OrniTela Inc., Lithuania). The Common Cranes were captured during their arrival at the roosting wetland and while foraging during daylight hours. In the roosting wetland, mist nets were strategically deployed at the appropriate angle to the direction of the Common Crane arrival during dusk, and in foraging fields, they were captured using slip-knot leg nooses. Upon capture, each Common Crane was immediately extracted and blindfolded to minimize stress. Subsequently, a 40 g L-40 transmitter was fitted to the tibia of each Common Crane, adding approximately 0.93% to the average weight of the tagged Common Cranes. The deployment of transmitters was completed within a minute, morphological measurements were taken and the tagged Common Cranes were released within 17-30 minutes after capture. All transmitters were programmed to record GPS locations at 10-minute intervals, which were then transmitted via a 2G GSM network.

### Data processing and migration parameters

The transmitter data was downloaded in CSV format and was subsequently visualized in QGIS 3.34.5 (52) to look for any errors in GPS locations, during which unusually deviated data points were removed. Trajectories between the breeding and wintering grounds were extracted in QGIS 3.34.5. To delineate summering and wintering areas, we combined spatial criteria with behavioral movement patterns derived from GPS tracking data. An individual Common Crane was considered to have arrived ("arrival date") at its seasonal destination (summering or wintering ground) when it entered a known range for that season and began exhibiting consistent localized movements-defined as daily displacements of less than 30 km/day around a fixed site-for 15 or more consecutive days. This movement threshold was determined based on the highest mean daily distances recorded from the Common Crane settled on their respective summering and wintering grounds. Given that the Common Crane repeatedly returns to the same roost site (53,54), these phases were characterized by typical looping or cyclical movements between foraging fields and roosting wetlands. The "departure date" was defined as the first day when the bird transitioned from this localized activity to sustained, directional long-distance movement(> 100 km in a day), leaving known seasonal ranges in subsequent days indicative of migratory travel. This marked the end of the stationary phase and the initiation of migration.

"Stopover days" were defined as days during which the bird remained within a 50 km radius during the day and showed low average travel speed (<10 km/h) and looping or localized movements, typically between roosting and foraging areas (53). The "migration duration" was calculated as the total time taken by a Common Crane to travel between the breeding and wintering grounds, including stopovers. Specifically, for autumn migration, we used the last GPS location recorded at a breeding site before departure and the first GPS location recorded at a wintering site upon arrival. The same approach was applied in reverse for spring migration. Although cranes often used multiple sites within the broader breeding or wintering range, these sites were typically clustered within a 100 km radius and formed part of a single seasonal area. The "travel duration" was defined as the total number of travel days during migration, subtracting any days spent at stopover sites. The "migration distance" was measured as the cumulative travel distance in km between all consecutive GPS locations during migration. The "migration speed" was defined as the total migration distance divided by the total duration of the migration, expressed in km/day. The "travel speed" was calculated as the daily distance traveled during the travel days, where the total migration distance was divided by the total travel duration and expressed as km/day.

The "stopover duration" was the aggregate of all days spent at various stopover sites during each migration season. Further, the "straightness index" was calculated to measure orientation efficiency, defined as the ratio of the straight-line distance (D) between the breeding and wintering sites (for autumn migration) or wintering and breeding sites (for spring migration) to the actual path length (L) traveled by the bird. For this, the straight lines were simulated for each migration trajectory and the ratio D/L was calculated to determine an empirical measure of the directness of the migration path. To assess the extent of longitudinal detours adopted by each individual, we computed the linear distance of each location along the actual migration path from the simulated straight line (55). The mean longitudinal detours were calculated for each latitude to pinpoint the latitudes exhibiting the highest degree of detour.

To examine seasonal variation in migration corridor, we first rounded all GPS locations to the nearest full-degree latitude. For each latitude bin and for each season (Autumn and Spring), we calculated the average longitude of all GPS fixes to define the population-level mean migratory axis. We then measured the great-circle distance between each GPS point and the corresponding mean longitude at its latitude and season, providing a measure of longitudinal deviation in kilometer. These values represent how widely birds travelled around the central route. We summarized the mean and confidence interval of longitudinal deviation by latitude and season, and applied a loess smoother to visualize how the width of the migration corridor varied along the latitudinal gradient in spring and autumn (56).

### High-use areas and migratory bottlenecks

To identify areas of high space use across the migratory population, we calculated dynamic Brownian Bridge Movement Models (dBBMMs) for each individual migration track using the package move (57) in R version 4.5.0 (58). To derive population-level patterns of use, all individual dBBMMs were stacked and summed pixel-wise to create a cumulative utilization surface. This surface was then normalized to obtain a population-level utilization distribution (UD), from which we extracted the 50%, 75%, and 99% isopleths. The 50% isopleth represents high-use areas, 75% isopleths represents moderate use areas while the 99% isopleths represents the migration range map of all the tracked Cranes.

To identify migratory bottlenecks, we used 99% UDs generated for each individual using dBBMM (59). Each UD was converted into a binary raster indicating presence (1) or absence (0) of use across the range. These binary layers were then summed across individuals to produce a cumulative raster representing the number of overlapping migration routes at each pixel. A 10 x 10 km grid was overlaid on the study area, and the mean overlap value within each grid cell was extracted from the cumulative raster. This mean was normalized by the total number of UDs to calculate the proportion (ranging o to 1) of individuals using each cell. Grid cells with a proportional overlap value of 0.8 or higher were interpreted as migratory bottlenecks, representing areas of pronounced spatial convergence across multiple migration trajectories.

### Statistical analysis

To assess the impact of migration season on key migration parameters, a General Linear Model (GLM) was performed, employing ordinary least squares (OLS) regression. Bootstrapped confidence intervals were generated using 1000 samples and the bias-corrected and accelerated (BCa) method to ensure accurate parameter estimates and address potential deviations from normality in the residuals. Heteroscedasticity-consistent standard errors (HC3) were applied to account for potential violations of homoscedasticity. The model fit was evaluated using R^2^• The summary statistics for each migration parameter were calculated and data is presented as mean ± SE and range. The visual comparison of different parameters between outbound and inbound migrations were presented using dot whisker plots where whiskers show 95% Cl.

To investigate habitat preferences of Common Cranes during migration stop-over periods, we implemented a Resource Selection Function (RSF) using logistic regression, operationalized as a binomial Generalized Linear Model (GLM). The analysis compared habitats at locations that the Cranes used during stop-over periods (termed "used") with habitats that were available but not used by the birds in the same space and time frame. To reduce spatial and temporal autocorrelation, we randomly selected 10% of the used GPS locations at each stop-over site for each tagged Crane. Around these used points, we generated a 5 km union buffer and sampled "available" (i.e., not-used) points at a tenfold ratio relative to the number of used points within this buffer. We extracted the habitat types for all points using 10 m resolution ESRI land use/land cover raster layers downloaded from https://livingatlas.arcgis.com/landcover/ for the year 2023.

Using the package lme4 (60) in R version 4.5.0, we initially fit a Generalized Linear Mixed Model (GLMM), including individual migration events as a random effect to account for potential variation among birds. However, the variance explained by the random effect was minimal (variance= 0.019, SD= 0.13), and the model exhibited signs ofoverfitting. We therefore compared model fit using Akaike Information Criterion (Ale) and Bayesian Information Criterion (BIC) values between the GLMM and GLM. The GLM showed better support (GLM AIC = 5645.4, BIC = 5688.2 vs. GLMM AIC = 5643.2, BIC = 5693.2), and was selected as the final model based on its lower complexity and more interpretable structure. Finally, the GLM was fitted with a binary response variable (used= 1, available= 0) and habitat type as a categorical predictor. The model estimated the relative probability of use across different habitat types at stop-over locations, using "Rangeland" as the baseline category. Predicted probabilities and confidence intervals were extracted and visualized to interpret habitat selection patterns.

## Results

A total of 40,336 GPS fixes were collected attributed to the migratory journeys of five Common Crane individuals across three years (Supplementary table S1). One of the individuals tagged in 2020 completed a full migration cycle (defined as complete Autumn and Spring migration) before data transmission ceased, while the other four individuals tagged in 2022 completed two full migration cycles each, resulting in a total nine complete migration cycles containing 18 trajectories.

### Migration timing, routes and stop-over sites

During their Spring migration, the birds left the wintering grounds for their breeding grounds between March 18 and April 11, with an average departure date of March 29. They arrived at the breeding grounds between April 9 and April 26, averaging an arrival date of April 19. For their return journey, the birds began migrating back to India from South-Western Siberia (55°N - 58°N) between September 28 and October 12, with an average departure date of October 5. They reached Gujarat, in Western India (22°N - 23°N), between October 6 and October 25, with an average arrival date of October 16.

The migration route of all tagged Common Cranes passed through Pakistan, Afghanistan, northwest Iran, Turkmenistan, Uzbekistan, and Kazakhstan, ending in southwestern Siberia and northern Kazakhstan (Figure 1 and 2). During both migrations, the cranes never crossed the Hindukush Himalayas. Along the route, they also traversed the vast Registan, Dasht-e-Margo, and Karakum deserts. On the Spring journey, the cranes initially flew in a northwest direction towards the Aral Sea at 45°N latitude. From there, they turned northeast to reach their breeding grounds. However, on two occasions, the cranes maintained a northwest direction until reaching S0°N before heading northeast. Notably, the cranes never crossed S6°E, the Western limit of the Ural Mountains of Russia.

**Figure 1.**
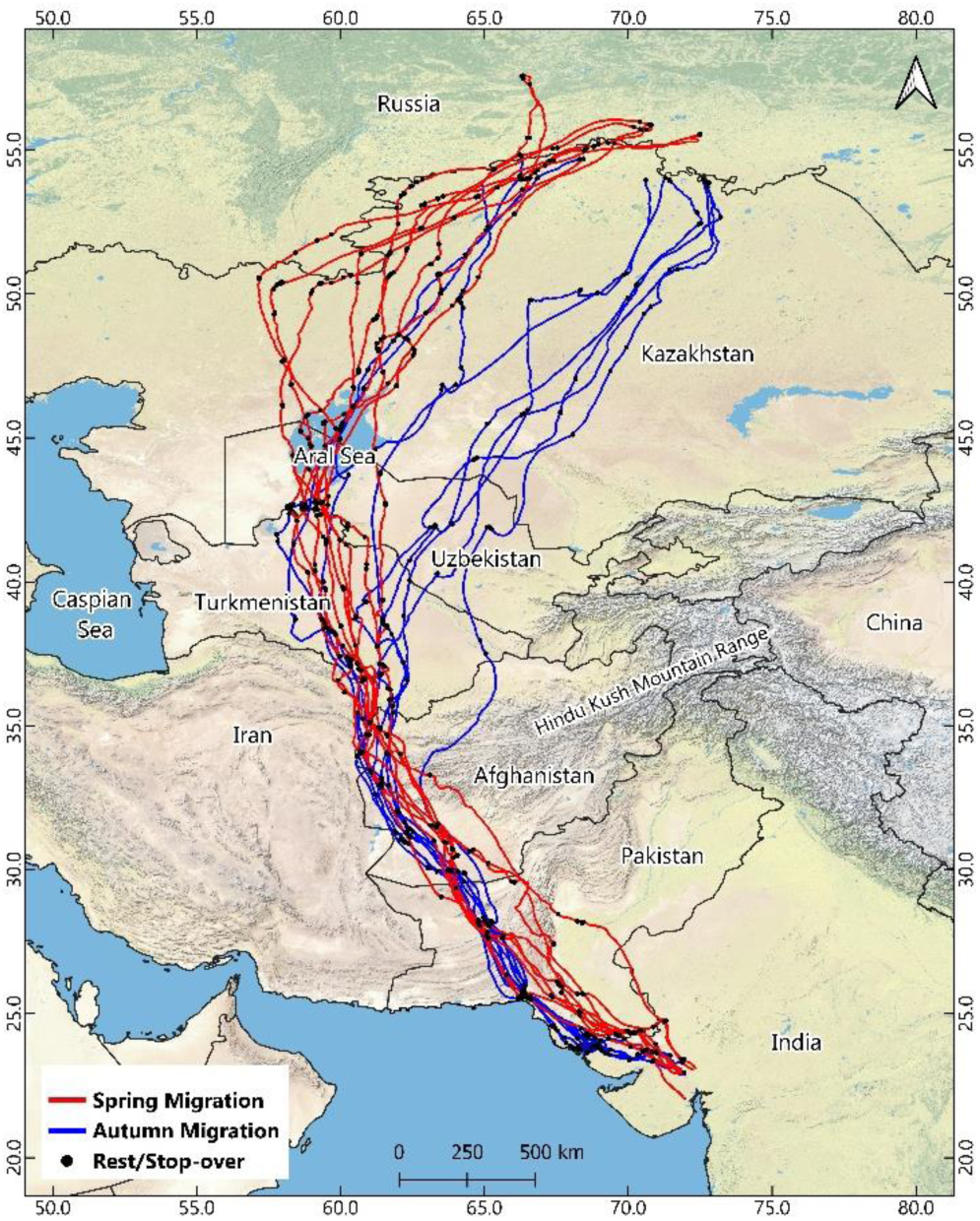
A map showing migration trajectories of five tagged Common Cranes between 2020 - 2023. (Base map: ESRI Physical)

**Figure 2.**
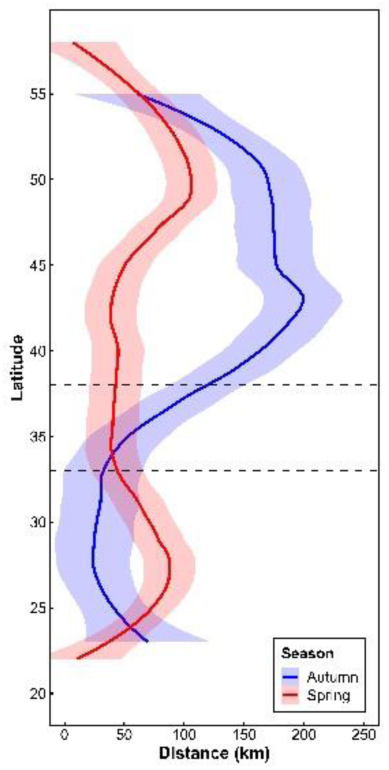
LOESS-smoothed Longitudinal variation (in kilometer) across latitude during Autumn and Spring migration. Solid lines represent the mean longitudinal deviation, and shaded areas indicate 95% confidence intervals. Dashed horizontal lines mark the latitudinal extent of the Hindu Kush mountains, a potential topographic barrier along the flyway.

Analysis of longitudinal deviation across latitude revealed clear seasonal differences in migration corridor alignment. In Spring, cranes followed a more westerly route that extended across the Aral Sea region, while Autumn migration took place along a more easterly pathway, with no indications of birds travelling near the Aral Sea. The two seasonal routes remained spatially distinct up to approximately 38°N latitude, after which they gradually converged. South of this latitude, both Spring and Autumn migrations show spatial overlap as they traverse Turkmenistan, Afghanistan, and Pakistan before reaching the wintering grounds in western India. These patterns indicate that migration routes differ between seasons in the northern part of the flyway but become increasingly aligned further south (Figure 2).

The dBBMM analysis revealed that high-use areas during migration were concentrated in four broad ecological regions: the arid deserts of Southern and Western areas of Central Asia, the riverine floodplain of the Amu Darya, Aral Sea and the open steppe landscapes of northern Kazakhstan and Southern Russia. Prominent desert stopover sites included the Rann of Kutch Ramsar Site and Kharan desert in Pakistan, north-western reaches of the Dasht-e-Margo desert of Afghanistan, in Northern Iran where the political borders of Iran, Afghanistan and Turkmenistan meet and the regions of the Karakum Desert near Ashgabat, Mary, and Kerpichli in Turkmenistan, as well as Tejen and the Hanhowuz Reservoir. In the steppe regions, high-use areas were located in Central and North Kazakhstan. (Figure 3).

**Figure 3.**
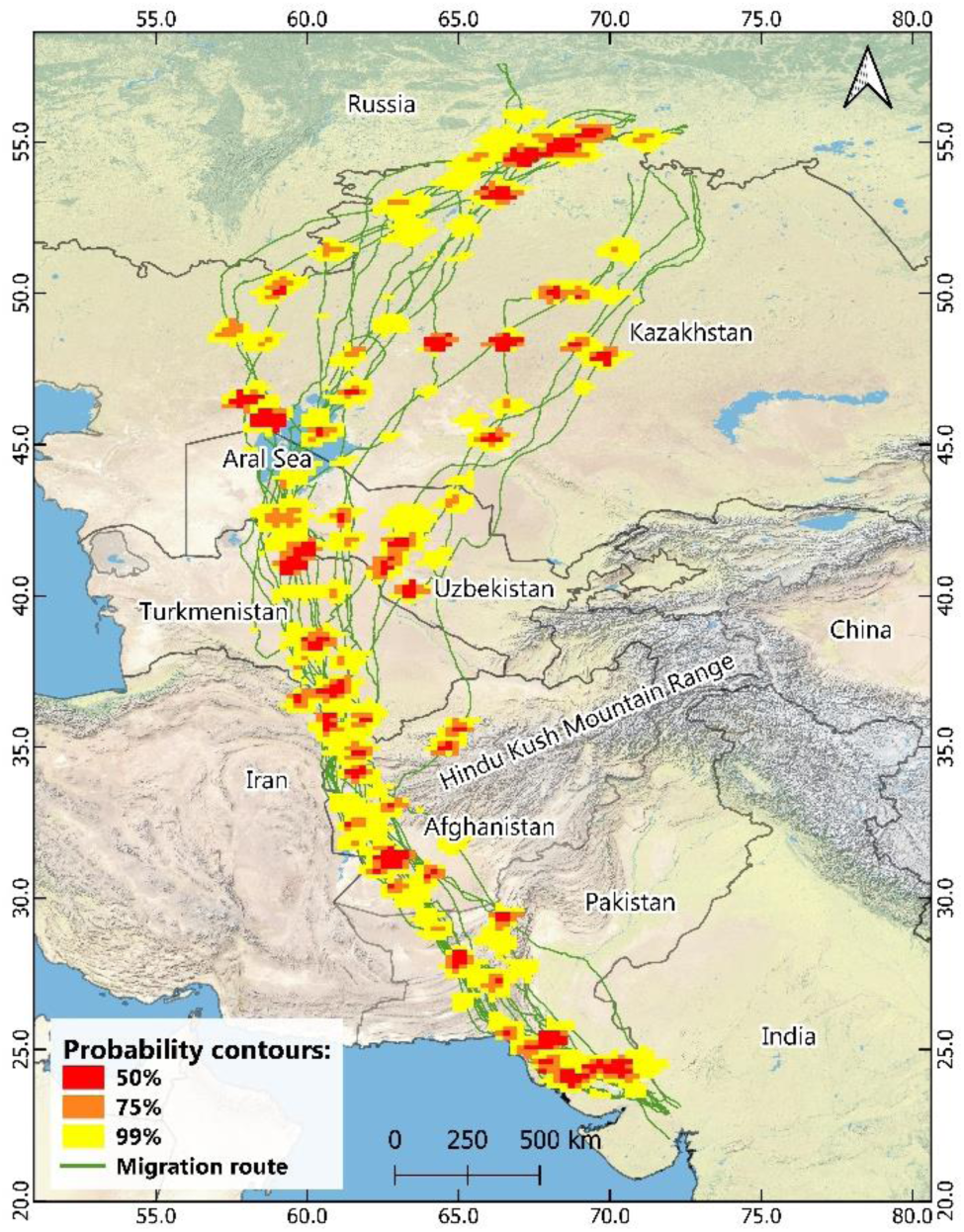
A map showing the population-level utilization distribution of the tracked Common Cranes derived through dynamic Brownian Bridge Movement Model. (Base map: ESRI Physical)

### Seasonal differences in migration parameters

The results of the General Linear Model (GLM) analyses highlight significant seasonal differences in migration parameters between Autumn and Spring migration directions. The model for migration distance exhibited a strong fit (R^2^ = 0.78, β^ = 652.67; SE = 90.91, 95% Cl = 490.30 - 817.78, p = <0.0001), indicating that Spring migration covers significantly longer distances. Similarly, the model for migration duration revealed a robust fit (R^2^ = 0.68, β^ = 10.11; SE= 1.86, 95% Cl= 7.05 - 13.72, p = <0.0001), reflecting a notable increase in migration duration for Spring migration.

For migration speed, the analysis indicated a strong fit and a significant difference (R^2^ = 0.64, _β^_ = 147.75, SE= 29.47, 95% Cl= -97.13--5.01, p = 0.00013) where the Spring migration was associated with a marked decrease in speed. The stopover duration also demonstrated a strong model fit (R^2^ = 0.61, _β^_ = 6.33, SE= 1.34, 95% Cl = 3.91 - 8.65, p = 0.00024), indicating increased resting periods during Spring migration. Furthermore, the analysis of the straightness index (SI) revealed a low model fit (R^2^ = 0.48, _β^_ = -0.05, SE= 0.01 95% Cl= -0.07 --0.02, p = 0.00231), suggesting that Spring migration exhibited a slight reduction in straightness, indicating less direct routes. The travel speed showed the weakest model fit and no significant difference between the Autumn and Spring migration (R^2^ = 0.20, β^ = -60.61, SE= 31.73, 95% Cl= -124.03-2.81, p = 0.07). The summary statistics of the key migration parameters are given in Table 1 and supplementary table S2, and are graphically compared in Figure 4.

**Figure 4.**
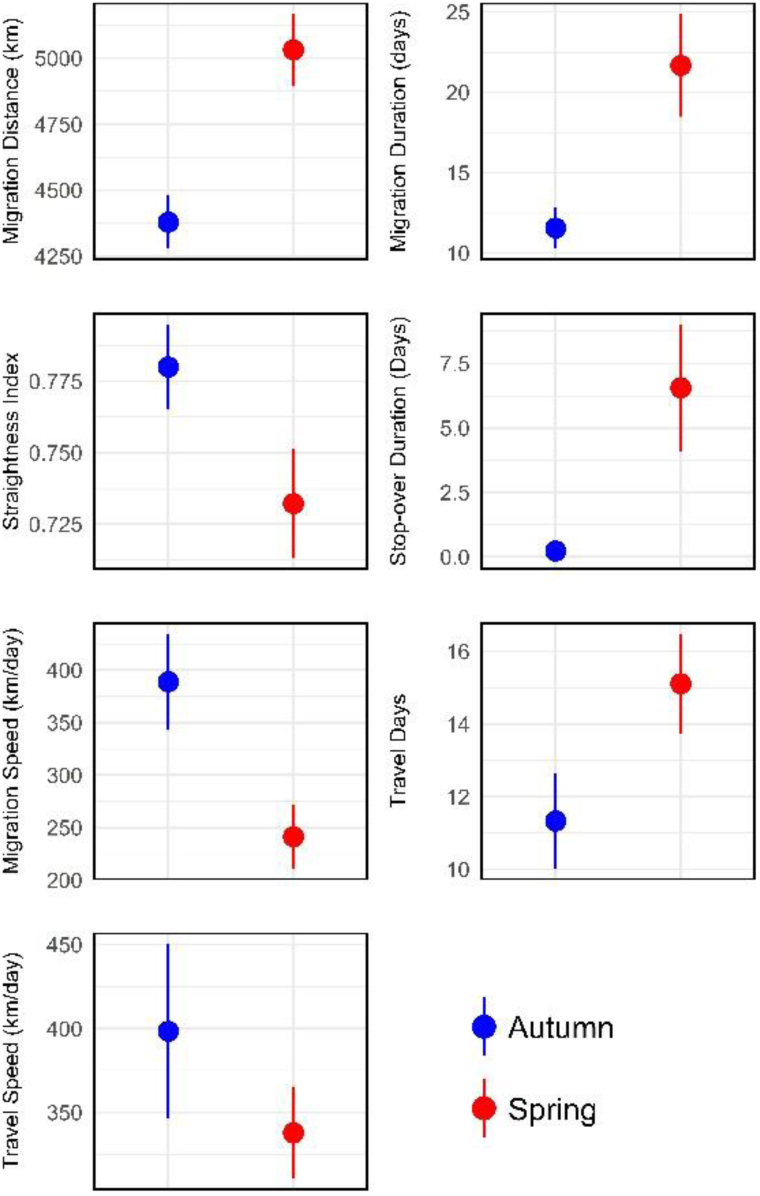
Dot-whisker plots comparing the key migration parameters of the five tagged Common Cranes. The dots show mean values and the whiskers show 95% confidence interval.

**Table 1.** A table showing the summary statistics of key migration parameters of the five tagged Cranes during Autumn and Spring migration journeys.

| Parameter | Migration Direction | Mean | Median | SD | Range |
| --- | --- | --- | --- | --- | --- |
| Migration Distance (km) | Autumn | 4378.8 | 4340.0 | 153.1 | 4165.0 - 4593.0 |
|  | Spring | 5031.4 | 5046.0 | 206.6 | 4638.0 - 5278.0 |
| Migration Duration (days) | Autumn | 11.6 | 12.0 | 1.9 | 9.0 - 14.0 |
|  | Spring | 21.7 | 22.0 | 4.9 | 15.0 - 31.0 |
| Straightness Index | Autumn | 0.8 | 0.8 | 0.0 | 0.7 - 0.8 |
|  | Spring | 0.7 | 0.7 | 0.0 | 0.7 - 0.8 |
| Stop-over Duration (days) | Autumn | 0.2 | 0.0 | 0.4 | 0.0 - 1.0 |
|  | Spring | 6.6 | 8.0 | 3.8 | 0.0 - 12.0 |
| Migration Speed (km/day) | Autumn | 388.9 | 381.2 | 69.5 | 310.0 - 500.8 |
|  | Spring | 241.2 | 239.9 | 46.1 | 169.0 - 320.5 |
| Travel Days | Autumn | 11.3 | 12.0 | 2.0 | 8.0 - 14.0 |
|  | Spring | 15.1 | 15.0 | 2.1 | 12.0 - 19.0 |
| Travel Speed (km/day) | Autumn | 398.4 | 381.2 | 79.6 | 324.1 - 551.4 |
|  | Spring | 337.8 | 330.9 | 41.4 | 275.8 - 394.7 |

### Migratory detour and bottleneck

During Autumn migration, detours commence from the highest latitude of 55°N, where minimal detour distances (11.44 km) are observed, and progressively increase as latitude decreases toward the southern latitudes and peaks at 34°N with the detour of 829.56 km. Beyond this peak, detour distances decrease at lower latitudes, as the trajectories approach wintering ground. During the Spring migration, the detour distances initially increase gradually, showing moderate detours from 22°N to 29°N. A more pronounced increase in detour distances is observed between latitudes 29°N and 35°N, with the highest detour distance of 969.32 km recorded at 35°N. Post-peak, Spring detours decline toward higher latitudes, as the trajectories approach the breeding ground (Figure 5). The observed detours resulted in migration routes that were 23.3% (SE= 1.29, 95% Cl = 25.3-31.2) longer in Autumn and 36.8% (SE= 1.85, 95% Cl= 32.5-41.0) longer in Spring compared to the shortest straight-line distance between breeding and wintering sites.

**Figure 5.**
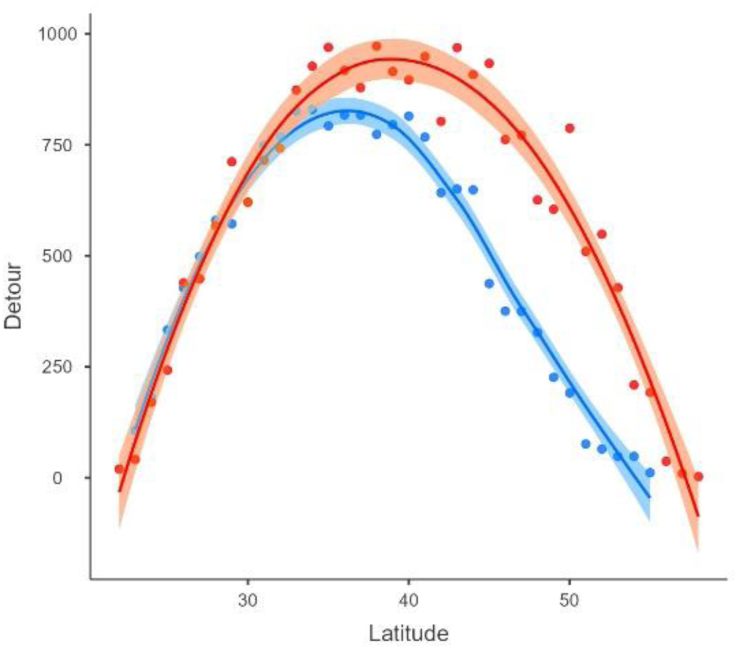
A graph showing the seasonally explicit spatial patterns in the migratory detour, visualised by fitting smoothed LOESS lines. Red line shows Spring migration, blue line show Autumn migration, shaded areas show 95% confidence interval. The values of detour are in km.

The 99% UD overlap analysis identified three distinct regions along the migration corridor where the proportional use ranged from 0.8 to 1.0, indicating that these grid cells were utilized by 80% or more of the tracked individuals. These regions were designated as migratory bottlenecks, representing zones of pronounced spatial convergence across multiple migration trajectories. The identified bottlenecks include: (i) Ahal province, Turkmenistan along the Tedzhen River near the Iran border, (ii) the North-Western reaches of the Dasht-e-Margo desert in Afghanistan, and (iii) the Kha ran Desert region of Pakistan (Figure 6).

**Figure 6.**
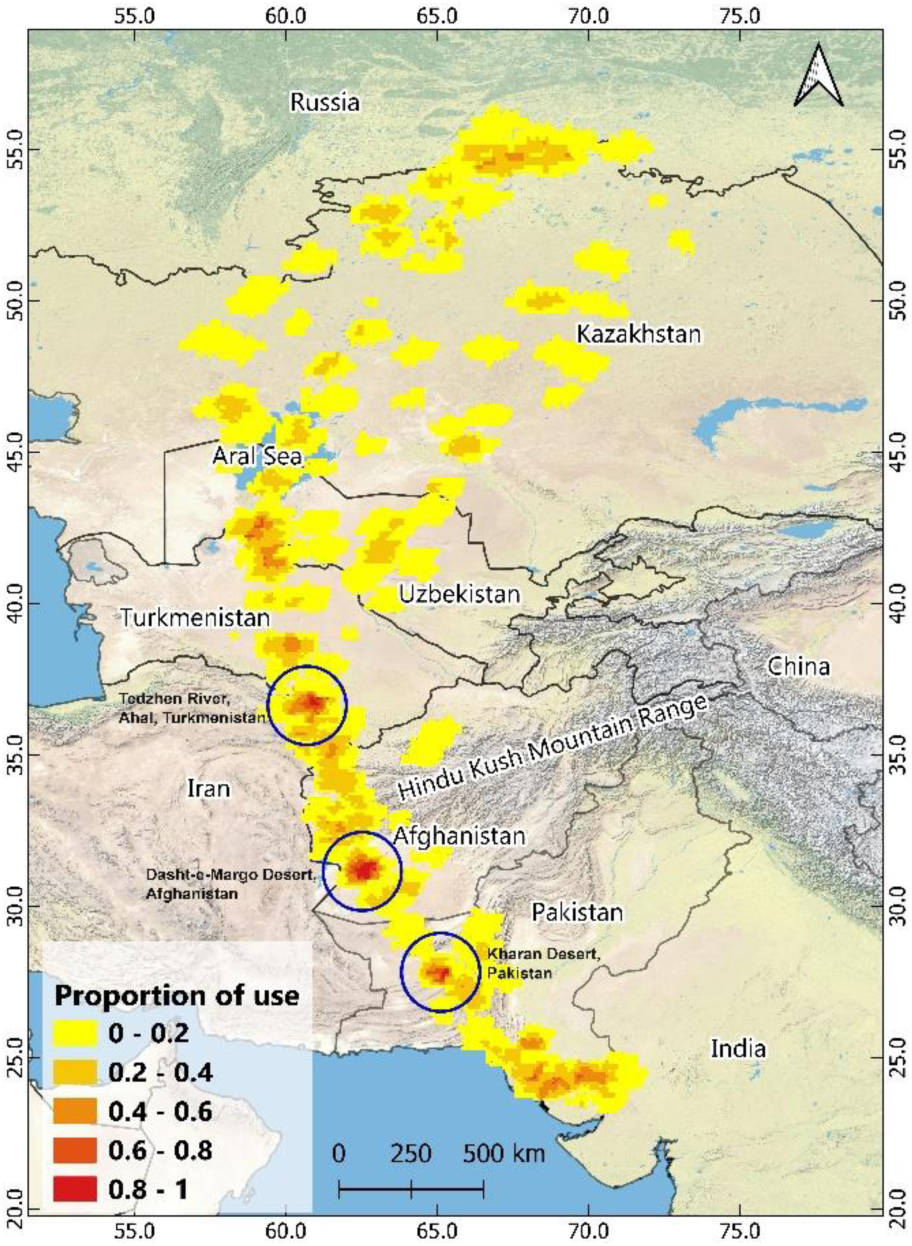
A figure showing three major migratory bottlenecks (areas with proportion of use more than 0.8) along the migratory route of the tracked Common Cranes_ The bottlenecks are indicated with blue circles_ (Base map: ESRI Physical)

### Habitat Selection at Stopover Sites

The resource selection function (RSF) analysis indicated varying probabilities of use across the available habitat types at stopover locations. Bare ground had the highest predicted probability of use (0.18; 95% Cl: 0.13-0.23), followed by cropland (0.13; 95% Cl: 0.11-0.14) and water bodies (0.09; 95% Cl: 0.06-0.14). Lower probabilities were observed for rangeland (0.07; 95% Cl: 0.07-0.08), flooded vegetation (0.06; 95% Cl: 0.04-0.11), and forest (0.02; 95% Cl: 0.00-0.11) (Figure 7).

**Figure 7.**
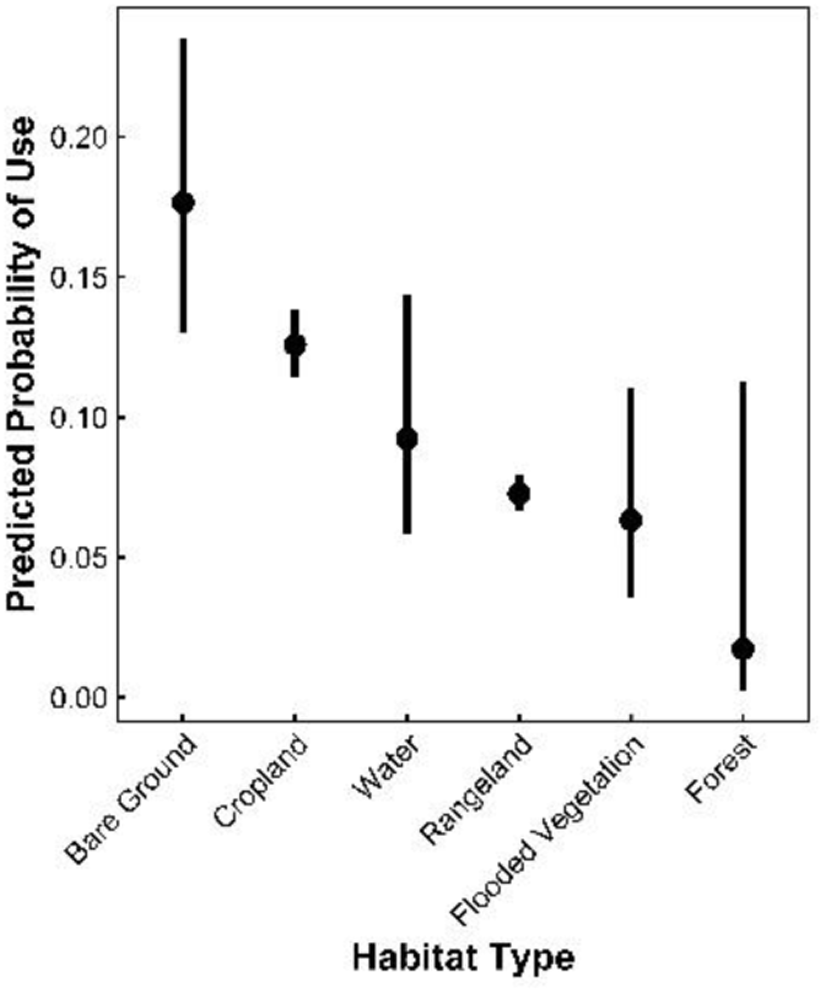
Dot-whisker plot showing predicted probabilities of habitat use by Common Cranes at stopover sites during migration. Dots represent mean predicted probabilities for each habitat type, and whiskers indicate 95% confidence intervals.

## Discussion

### Migration timing, routes and stop-over sites

The migration timings and routes observed in our study, based on multi-year tracking of five tagged Common Cranes, offer a more comprehensive understanding compared to previous studies, which tracked fewer Common Cranes over a single season or year. The Spring migration timing aligns with prior research by (41,42) and (43), where cranes departed Western India in March and early April. The Autumn departure date is consistent with one of the two individuals tracked in (43) where the Crane departed from Russia in the last week of September. However, the second individual from (43) departed on 14th August, which is 52 days earlier than the mean departure date recorded in our study, suggesting the individual variability in departure decision. The migration timings recorded in our study establish a crucial baseline for Common Cranes along the Central Asian Flyway, providing a foundation for detecting climate-driven phenological shifts. Long-term studies in Europe and Russia have demonstrated significant advances in the Common Crane migration timing in response to rising temperatures (61,62), suggesting the role of climate change in altering avian migration phenology. Given the increasing impacts of global warming, continued monitoring of migration phenology is essential for understanding species-level responses and informing conservation strategies.

The broad migration route crossing Pakistan, Afghanistan, Northern Iran, Turkmenistan, Uzbekistan and finally reaching the breeding ground in South-Western Siberia is consistent with (42,43). While the breeding region of the cranes is also consistent with (41), the Spring migration route differs slightly, as their Common Cranes did not go all the way to Northern Iran and the Aral Sea and flew through Northern Afghanistan instead of Southern Afghanistan. This variation stems from the Northern Indian origin of their Crane population. The dual migration corridors observed in our study-western during Spring and eastern during Autumn-likely reflect seasonal shifts in environmental conditions. Such seasonal routing is frequently driven by variation in resource availability and environmental patterns. For example, (63) demonstrated that Barnacle Geese *Branta /eucopsis* adjust their Spring migration to follow the "green wave" of vegetation growth in the Western Palearctic, while (11) found that honey buzzards on the East Atlantic Flyway alter their routes to exploit supportive wind regimes. In our case, the Common Cranes appear to detour westward during Spring to leverage better stopovers, whereas the Autumn route shifts eastward as the Common Crane do not make stopovers during the post-breeding return migration, likely prioritizing a more direct route when immediate energy accumulation is less critical. This major migratory corridor encompassing Spring and Autumn corridors used by the Common Crane in our study, as well as documented in previous research (42,43), highlights strong migratory connectivity between Western India and the breeding grounds in Northern Kazakhstan and South-Western Siberia. We refer to this major corridor as "the Desert Corridor" of the Central Asian Flyway to emphasize its ecological distinctiveness and consistent geographic structure. This route contrasts with other CAF migration corridors, which necessitate the Himalayan crossings for example by the Demoiselle Crane (64–66), Raptors (57–59), and Goose and Duck species (70,71). Recognizing such sub-corridors is important for improving ecological understanding and conservation planning, especially in regions like this, where migration data remain scarce compared to better-studied routes. Defining the Desert Route helps prioritize key habitats and bottlenecks, and informs region-specific strategies for safeguarding long-distance migrants along this underrepresented portion of the CAF. The migration distances recorded in our study are consistent with those reported by (42) and one individual tracked by (43). However, another individual in (43) was reported traveling 12,982 km during the Spring migration and 14,873 km during the Autumn journey. These distances are 2.5 times (Spring) and 3.3 times (Autumn) greater than those recorded in our multi-year study. Given the similarity in breeding and wintering sites between studies, this discrepancy is likely due to differences in analytical approaches.

The cumulative RSF-based habitat selection analysis, performed across all identified stopover sites, indicated the highest predicted use for bare ground, followed by cropland and water bodies. This suggests that during migration, Common Cranes tend to select open, resource-accessible landscapes that may provide both foraging opportunities and enhanced visibility, important for minimizing predation risk and maintaining social cohesion. While the analysis did not evaluate habitat selection separately for each stopover site, the site-level data contributed cumulatively, allowing us to identify consistent preferences across the migration corridor. Notably, several key stopover regions-including the Karakum Desert, Amu Darya floodplains, and northern Kazakh steppe-are dominated by the land cover types associated with higher predicted use. The Karakum Desert, including sites near Ashgabat, Mary, Kerpichli, and Tedzhen, serves as a crucial staging area despite its arid conditions, emphasizing the species’ ability to exploit ephemeral water sources and localized productivity within desert ecosystems (72). Similarly, the Amu Darya floodplains function as essential stopover sites, offering stable wetland as well as food resources in the form of food crops in an otherwise arid landscape, reinforcing the well-documented importance of riparian corridors in Crane migration (73). Steppe habitats in Northern Kazakhstan, including the Aktobe and Kostanay regions, as well as in North Kazakhstan, represent key staging areas that likely support extensive refuelling due to the availability of agricultural fields and natural grasslands. The prolonged use of these sites aligns with previous studies demonstrating that Common Cranes frequently exploit human-modified landscapes during migration (74–76). Furthermore, the inclusion of Adamovsky district, Oren burg Oblast (Russia), and Taybad (Iran) as stop-over locations indicates the species’ reliance on a broad network of habitat patches to meet the energetic demands of long-distance migration.

Given the increasing anthropogenic pressures on natural habitats within the Central Asian Flyway (26), the conservation of these critical stop-over sites requires a multi-scalar approach integrating habitat protection, sustainable land-use practices, and transboundary conservation initiatives. Further, the identified stopover sites likely host high densities of migratory birds, creating potential hotspots for disease transmission. Recent outbreaks of highly pathogenic avian influenza (HSN1) have caused mass mortality in Common Cranes (77,78), highlighting the vulnerability of migratory populations to emerging infectious diseases. Given the risks associated with high-density aggregations, regular disease monitoring and screening at key stopover sites are essential for early detection and mitigation. Integrating such measures into transboundary flyway conservation and management plans will be critical for safeguarding both migratory bird populations and ecosystem health.

### Seasonally explicit migration duration

The analysis of migration parameters revealed that Spring migration in Common Cranes was significantly slower than Autumn migration, with longer stopovers contributing to the extended duration. This contrasts with findings from the Eastern Mongolian Common Crane population (76), where Spring migration was faster. which was also true many other long-distance migratory species (79), however, these species are predominantly small in size. Faster Spring migration is typically linked to securing better territories, enhanced reproductive success, and improved readiness for subsequent migrations (80–82). Like in our case, the slower Spring migration is also reported in many other long-distance migrants with large body sizes such as the White Stork *Ciconia ciconia* (83), the Common Eider *Somateria mollissima* (84), the Black-backed Gull *Larus fuscus fuscus* (85), the Tundra Swan (Bewick’s) *Cygnus columbianus bewickii* (86), the Booted Eagle *Hieraaetus pennatus* (87), the Egyptian Vulture *Neophron percnopterus* (88) and the Greater White-fronted Goose *Anser a/bifrons* (16, 66). These patterns suggest that seasonal variation in migration duration and departure decisions is likely influenced by an interplay of factors including body size, the extent and timing of pre-migration fuel accumulation and energy expenditure strategies, migration distance, flight mode (e.g., soaring vs. flapping), and the spatial distribution of stopover opportunities. Thus, rather than a simple dichotomy, migration timing reflects a complex trade-off between ecological conditions, physiological constraints, and life-history priorities.

Given that cranes are long-lived capital-income breeders, the slower Spring migration observed in our study individuals may reflect the need for females to accumulate energy reserves for reproduction-a strategy also documented in geese, another large-bodied grazing bird (89,90). Additionally, the Common Crane may delay migration to coincide with snowmelt at their northern breeding grounds, ensuring that breeding areas are accessible and favorable upon arrival (86,91). This behavior helps the Common Crane optimize their reproductive success, as arriving too early could mean facing harsh conditions on breeding grounds. The proximate factors governing stopover duration and departure decisions during Spring migration by the Common Crane in the CAF warrant further investigation.

### Migratory detour and bottleneck

This study presents the first quantitative assessment of detours undertaken by Common cranes along the CAF, revealing a substantial deviation from the shortest possible migration route. Earlier, migratory detours have been examined in White-naped Crane *Grus vipio* where the Cranes took detours to capitalise on high-quality habitat (47). Conversely, the Black-necked Crane *Grus nigricollis,* a crane species native to Asia, did not exhibit detours while migrating within the Tibetan Plateau (92). A more relevant comparison can be drawn with the Demoiselle Crane *Anthropoides virgo,* which shares wintering grounds with the Common Crane in Western India. The Demoiselle Crane breeding in northern China and Mongolia exhibit a pronounced loop migration: they cross the Himalayas during autumn but detour around the western Himalayas in Spring in response to seasonal variation in wind support, temperature, and resource availability (64,66). While loop migration involves seasonally distinct routes, the Common Cranes in our study followed detours within both seasonal migrations, rather than forming a loop. Spring detour in the Demoiselle Crane, reduced energetic costs by exploiting more favorable habitats and avoiding the Himalaya. Thus, both species appear to respond to similar ecological constraints along the CAF, but adopt different spatial strategies to optimize migration. As the long-distance migration is inherently constrained by the energetic demands of sustained flight (93), migratory birds frequently exhibit route adjustments to optimize refuelling opportunities and minimize physiological stress and the risk of predation (12,94,95). Our findings demonstrate that detours increased the total migration distance by up to 41% compared to the shortest route. The significant increase in migration length observed across latitudes during both spring and autumn migrations suggests that environmental and topographical factors strongly influence route selection.

The shortest straight migratory trajectory would necessitate crossing the Hindu Kush and the Kashmir Himalaya. The complex topography of these high-altitude regions is characterized by turbulent wind regimes, low atmospheric pressure, and unpredictable thermal availability (64,96), conditions that are largely unsuitable for facultative soaring species like the Common Crane. Given that the Common Crane also relies on thermal uplift to sustain long-distance flights with minimal energy expenditure, the scarcity of stable thermals in this region likely renders the shortest route energetically expensive. Consequently, the observed detours reflect a strategic avoidance of suboptimal flight conditions, favoring alternative routes that facilitate soaring flight and reduce reliance on energetically costly flapping flight.

The westward deviation observed in both Spring and Autumn migrations directs the cranes through the arid landscapes of Pakistan, Afghanistan, and Turkmenistan, where extensive desert expanses generate strong and consistent thermal uplift and also provide wind assistance (64,97). The availability of such atmospheric conditions likely enables Common Cranes to employ a mixed flying strategy, mitigating the energetic costs associated with increased detour distances. Beyond the benefits conferred by thermal availability, the presence of critical stopover habitats along this route appears to further justify the observed deviations from the shortest path. The agricultural landscapes near the Iran-Afghanistan border, wetland complexes and desert oases in Turkmenistan, and the highly productive agricultural fields of the Amu Darya River Valley provide key resting and foraging opportunities, enabling the cranes to replenish energy reserves at strategically spaced intervals. The pronounced detours and prolonged stopover period observed during Spring migration further highlight the role of habitat quality in influencing route selection, as cranes must accumulate substantial energy reserves before continuing into breeding grounds. By integrating thermal soaring opportunities with access to high-quality stopover habitats, the observed flyway effectively offsets the energetic costs associated with increased travel distance.

The identified migratory bottleneck at Southern Turkmenistan along the Tedzhen river close to the Iran border, north-western reaches of the Dasht-e-Margo desert of Afghanistan, and the Kharan desert of Pakistan highlights a critical conservation priority for the Common Crane. This bottleneck possibly results from geographic constraints such as the Aladagh, Binalud and East Iranian Mountain ranges in the West and the high reaches of the Hindu Kush mountains of Afghanistan in the East, forcing cranes to funnel through a narrow corridor. Such natural barriers shape migratory routes by limiting alternative flight paths, concentrating individuals into specific regions (51). However, this convergence makes the species highly vulnerable to habitat degradation, land-use changes, and direct threats such as hunting even if the birds are not resting or foraging in these areas (98). The consequences of migratory bottlenecks extend beyond localized threats. Because such areas channel a significant portion of the population of a species, disturbances can have disproportionate impacts on overall population dynamics (50). Recognizing bottlenecks as ecological priorities can help integrate their protection into broader conservation strategies, benefiting not only cranes but also other migratory species that rely on these corridors. By addressing threats at these critical junctures, conservationists can enhance the resilience of migratory populations and maintain the ecological integrity of these essential flyways.

While these findings offer important and novel insights into migration strategies and conservation priorities of the Common Crane, they should be interpreted in light of the study’s limitations. In particular, the relatively small sample size and limited geographic scope of tracking necessitate caution when generalizing to the entire CAF population. Despite the modest number of individuals tracked in this study (n = 5), we obtained nine complete migration trajectories, with four individuals tracked over two full migration cycles and one over a single cycle. We acknowledge that such sample sizes constrain the generalizability of behavioral inferences. However, the ecological relevance of the dataset is enhanced by the fact that all individuals were tagged at two wetlands in Western India that support approximately 30,000 Common Cranes-nearly 30% of the species’ wintering population in the region. This increases the likelihood that the tracked birds are representative of a significant proportion of the population using the CAF to migrate to Western India. While the patterns observed in migration routes, stopover regions, and bottleneck areas offer valuable ecological insights, we stress the need for future studies with larger sample sizes to more comprehensively capture population-level variation and strengthen conservation planning across the flyway.

## Conclusion

Our tracking study of nine migration cycles of five Common Cranes across three years provides critical insights into the migration timing, routes, stopover areas and stopover habitat selection in the Central Asian Flyway. Notably, this study is the first quantitative assessment of migratory detours and the identification of a migratory bottleneck of the Common Crane along the CAF. The findings reveal that the Common Crane deviate substantially from the shortest possible migration route, increasing their travel distance by up to 41% to optimize energy expenditure and access high-quality stopover habitats. This detour behavior is likely driven by the avoidance of physiologically demanding ecological barriers, the Hindu Kush and the Kashmir Himalaya, and the strategic use of thermal soaring conditions in arid landscapes. Additionally, the identification of migratory bottlenecks where geographic constraints funnel all the Crane migration routes, highlights a critical vulnerability in the migratory flyway of the Common Crane.

The study also reinforces the importance of long-term, multi-individual tracking in capturing the full spectrum of migratory behavior. The observed temporal consistency in migration schedules, coupled with individual variability in departure decisions, underlines the dynamic nature of the Common Crane migration. The identification of dual migration corridors-Western during Spring migration and Eastern during return migration-further demonstrates seasonal flexibility in route selection, likely driven by environmental and topographical constraints and breeding location. Furthermore, the reliance on arid landscapes, river floodplains, and steppe habitats for stopover sites highlights the ecological importance of these regions in sustaining migratory populations.

## Conservation implications

Key stopover habitats and migratory bottlenecks in Turkmenistan, Afghanistan, and Pakistan have crucial conservation implications for the Common Crane and other species using the Central Asian Flyway. The Common Crane used open natural ecosystems (ONEs) as Stopover sites, such as the Karakum Desert, Amu Darya floodplains, and Central and Northern Kazakhstan’s steppe habitats provide essential resources for refueling and resting. Anthropogenic modifications in the habitats in these areas could have cascading effects on the migratory population of not only the Common Crane but also other species using the same flyway, necessitating habitat protection and integration into transboundary conservation initiatives. Our findings reveal major migratory bottlenecks for Common Cranes Southern Turkmenistan along the Tejen river close to the Iran border, north-western reaches of the Dasht-e-Margo desert of Afghanistan, and the Kharan desert of Pakistan. The bottlenecks in Pakistan and Afghanistan overlap with areas of intensive hunting and illegal trade, where thousands of Demoiselle and Common Cranes are captured and bred in captivity under poor conditions, and where commercial trade is largely unregulated and illegal (27, 28). The concentration of migrating individuals in this region makes the population highly vulnerable to even localized threats, highlighting the urgent need for targeted conservation actions, including international cooperation, enforcement of anti-poaching laws, and awareness campaigns in key regions of Pakistan and Afghanistan.

Beyond habitat and hunting-related threats, disease outbreaks pose an increasing risk to migratory birds. The recent mass mortality of Common cranes due to highly pathogenic avian influenza (HSN1) highlights the vulnerability of birds congregating in large numbers. Given that the stopover sites identified in this study likely host a high abundance of migratory birds, they could serve as hotspots for disease transmission. Regular monitoring and screening at these key locations are essential for early detection and mitigation of emerging infectious diseases. Integrating disease surveillance into conservation strategies will be crucial for maintaining population health and ensuring the stability of the broader migratory network.

Importantly, the Common Crane can serve as a sentinel species for monitoring the ecological integrity of the Central Asian Flyway, particularly along the desert route spanning Pakistan, Afghanistan, Turkmenistan, Uzbekistan, and Kazakhstan. Its demonstrated reliance on a mosaic of habitats-including arid deserts, river floodplains, and arid and steppe croplands -highlights its ecological versatility and the importance of maintaining habitat heterogeneity across the flyway. The identification of key stopover regions, along with documented migratory detours and bottlenecks, emphasizes the species’ sensitivity to habitat availability and landscape continuity. These traits, combined with its wide distribution and long-distance migrations, make the Common Crane an ideal indicator for assessing flyway health and the impacts of environmental change. By prioritizing its conservation, we can safeguard not only this species but also the broader ecological network dependent on this vital corridor.

## List of abbreviations

CAF: Central Asian Flyway
dBBMM: Dynamic Brownian Bridge Movement Model
GLM: Generalised Linear Model
RSF: Resource Selection Function

## Declarations

### Ethics approval and consent to participate

We received permits to capture and tag Common Cranes from the office of the Principal Chief Conservator of Forest, Gujarat State Forest Department, vide letters WLP/ACF/RTC-3/C/1050-51/2018-19, dated 04-02-2019, and WL/RTC/28/C/852-853/2021-22, dated 19-01-2022.

### Consent for publication

Not applicable

### Availability of data and materials

The datasets used and/or analysed during the current study are available from the corresponding author on reasonable request.

### Competing interests

The authors declare that they have no competing interests.

### Funding

This research was funded by the PowerGrid Corporation of India Ltd. under its Corporate Social Responsibility initiative. The funding agency had no role in study design, data collection, analysis, interpretation, or conclusions.

### Authors’ contributions

HLB, RSK, and AB conceived the study. RSK and AB secured funding. HLB, RSK and GS arranged fieldwork logistics. HLB, GS and RSK participated in fieldwork. HLB conducted data analyses. HLB wrote the original draft of the manuscript. All authors reviewed, edited and approved the final manuscript.

## Acknowledgements

We thank the Director and Dean of the Wildlife Institute of India for their support during the study. We thank the Mr. Shwetank Pandit (Rtd. IFS) and and Mr. V.J. Rana (Rtd. IFS), Sh. P. Purushothama (IFS), Dr. Chaudhary (IFS), Mr. Dipak Solanki (GFS), Mr. Swapnil Patel (GFS) of Gujarat State Forest Department for their support in conducting fieldwork. We thank Mr. Gani Sama and late Mr. Karshan Padhar, the frontline staff of the Nal Sarovar Ramsar Site, for their expert help in trapping birds. We thank interns and field support staff for their support in field work. We thank Mr. Shailesh Thakor, caretaker at Thal Ramsar Site and the people of Vadla village for providing logistics while field work. Mention of trade names or commercial products in this publication is solely to provide specific information and does not imply recommendation or endorsement by the authors or the affiliated institution.

## Notes

### Competing Interest Statement

The authors have declared no competing interest.

### Summary of Updates

Methodological approach to identify high use areas and spatial bottlenecks along the migratory route. Added habitat selection at stopover sites.

