## Supplementary material for "Tracking the long way around: seasonal migration strategies, detours and spatial bottlenecks in Common Cranes wintering in Western India": Table S1

Table S1: A table showing basic information of tagged Common Cranes

| Sr. No. | Name of the individual | Date of tagging | Capture and Tagging location | Weight (kg) | No. of locations used in the study |
|---|---|---|---|---|---|
| 1 | Vadla | 12-03-2020 | Vadla Wetland (22.922671 N; 71.958432 E) | 4.7 | 3649 |
| 2 | Gani | 10-02-2022 | Thol Ramsar Site (23.139794 N; 72.405048 E) | 4.3 | 8283 |
| 3 | Sanand | 22-02-2022 | Thol Ramsar Site (23.135603 N; 72.401336 E) | 4.5 | 7759 |
| 4 | Nal | 04-03-2022 | Vadla Wetland (22.918991 N; 71.96749 E) | 4.7 | 9289 |
| 5 | Bhal | 05-03-2022 | Vadla Wetland (22.910345 N; 71.984059 E) | 4.6 | 11356 |

Table S2: A table showing the migration parameters and migration timings of the tagged Common Cranes

| Season | Individual | Migration Distance (km) | Migration Duration (days) | Straightness Index | Stopover Duration (Days) | Migration Speed (km/day) | Travel Days | Travel Speed (km/day) | Departure Date | Arrival Date |
|---|---|---|---|---|---|---|---|---|---|---|
| Autumn | Bhal_1 | 4593.00 | 14.00 | 0.81 | 0.00 | 328.07 | 14.00 | 328.07 | 28-09-2022 | 11-10-2022 |
| | Bhal_2 | 4340.00 | 14.00 | 0.80 | 1.00 | 310.00 | 13.00 | 333.85 | 12-10-2023 | 25-10-2023 |
| | Gani_1 | 4295.00 | 11.00 | 0.80 | 0.00 | 390.45 | 11.00 | 390.45 | 29-09-2022 | 09-10-2022 |
| | Gani_2 | 4165.00 | 10.00 | 0.78 | 0.00 | 416.50 | 10.00 | 416.50 | 11-10-2023 | 20-10-2023 |
| | Nal_1 | 4507.00 | 9.00 | 0.78 | 0.00 | 500.78 | 9.00 | 500.78 | 28-09-2022 | 06-10-2022 |
| | Nal_2 | 4214.00 | 13.00 | 0.79 | 0.00 | 324.15 | 13.00 | 324.15 | 11-10-2023 | 23-10-2023 |
| | Sanand_1 | 4574.00 | 12.00 | 0.76 | 0.00 | 381.17 | 12.00 | 381.17 | 05-10-2022 | 16-10-2022 |
| | Sanand_2 | 4411.00 | 9.00 | 0.76 | 1.00 | 490.11 | 8.00 | 551.38 | 11-10-2023 | 19-10-2023 |
| | Vadla | 4310.00 | 12.00 | 0.74 | 0.00 | 359.17 | 12.00 | 359.17 | 29-09-2020 | 10-10-2020 |
| Spring | Bhal_1 | 5046.00 | 26.00 | 0.78 | 10.00 | 194.08 | 16.00 | 315.38 | 31-03-2022 | 25-04-2022 |
| | Bhal_2 | 5240.00 | 31.00 | 0.77 | 12.00 | 169.03 | 19.00 | 275.79 | 27-03-2023 | 26-04-2023 |
| | Gani_1 | 5016.00 | 19.00 | 0.74 | 2.00 | 264.00 | 17.00 | 295.06 | 30-03-2022 | 17-04-2022 |
| | Gani_2 | 5278.00 | 22.00 | 0.74 | 8.00 | 239.91 | 14.00 | 377.00 | 31-03-2023 | 21-04-2023 |
| | Nal_1 | 4963.00 | 23.00 | 0.73 | 8.00 | 215.78 | 15.00 | 330.87 | 30-03-2022 | 21-04-2022 |
| | Nal_2 | 5163.00 | 23.00 | 0.72 | 8.00 | 224.48 | 15.00 | 344.20 | 18-03-2023 | 09-04-2023 |
| | Sanand_1 | 4638.00 | 17.00 | 0.72 | 5.00 | 272.82 | 12.00 | 386.50 | 30-03-2022 | 15-04-2022 |
| | Sanand_2 | 5131.00 | 19.00 | 0.70 | 6.00 | 270.05 | 13.00 | 394.69 | 25-03-2023 | 12-04-2023 |
| | Vadla | 4808.00 | 15.00 | 0.69 | 0.00 | 320.53 | 15.00 | 320.53 | 10-04-2020 | 24-04-2020 |
